# CCR9 signal termination is governed by an arrestin-independent phosphorylation mechanism

**DOI:** 10.1101/2024.11.01.621505

**Authors:** Thomas D. Lamme, Martine J. Smit, Christopher T. Schafer

## Abstract

The chemokine receptor CCR9 coordinates immune cell migration from the thymus to the small intestine along gradients of CCL25. Receptor dysregulation is associated with a variety of inflammatory bowel diseases such as Crohn’s and ulcerative colitis, while aberrant CCR9 overexpression correlates with tumor metastasis. Despite being an attractive therapeutic target, attempts to clinically antagonize CCR9 have been unsuccessful. This highlights the need for a deeper understanding of its specific regulatory mechanisms and signaling pathways. CCR9 is a G protein-coupled receptor (GPCR) and activates G_i_ and G_q_ pathways. Unexpectedly, live-cell BRET assays reveal only limited G protein activation and signaling is rapidly terminated. Truncating the receptor C-terminus significantly enhanced G protein coupling, highlighting the regulatory role of this domain. Signal suppression was not due to canonical arrestin-coordinated desensitization. Rather, removal of GPCR kinase (GRK) phosphorylation led to sustained and robust G protein activation by CCR9. Using site-directed mutagenesis, we identified specific phosphorylation patterns that attenuate G protein coupling. Receptor internalization does not correlate with G protein activation capabilities. Instead, CCR9 phosphorylation appeared to directly destabilize the interaction of G protein heterotrimers with the receptor. This interference could lead to rapid loss of productive coupling and downstream signaling as phosphorylation would effectively render the receptor incapable of G protein coupling. An arrestin-independent, phosphorylation-driven deactivation mechanism could complement arrestin-dependent regulation of other GPCRs and have consequences for therapeutically targeting these receptors.

## Introduction

Chemokine receptors are class A G protein-coupled receptors (GPCRs) that drive cell migration during immune homeostasis, inflammation, and development in response to small protein agonists called chemokines. Binding of chemokines activates the receptors, inducing key conformational changes that promote G protein heterotrimer coupling, which activates the G protein and leads to guanyl nucleotide exchange within the Gα subunit and separation from the Gβγ dimer. Chemokine-directed cell migration is driven by signaling cascades initiated by the dissociated, active G protein. The activated receptors also recruit GPCR kinases (GRKs) which initiate signal termination by phosphorylating serine and threonine residues on the receptor cytoplasmic face and C-terminus. These modifications lead to arrestin recruitment which canonically blocks further receptor signaling by sterically occluding the G protein coupling interface. Arrestins also coordinate receptor internalization from the plasma membrane, where the GPCRs are either degraded or recycled for another round of signaling. Together, these effectors rapidly translate stimulation into larger cellular effects, while regulating signaling and over activation [1,2].

The C-C chemokine receptor type 9 (CCR9) mediates T-cell maturation and migration from the thymus to the small intestine along gradients of its sole endogenous chemokine ligand CCL25 [3,4]. Upon activation, the receptor couples to G_i_ and G_q_ protein. These initiate the signaling cascades that mediate migratory responses. The specifics of CCR9 signal regulation are unknown, however, the activated receptors recruit arrestins and show desensitization following overstimulation [5]. Thus, signal termination can be assumed to follow the classical GRK to arrestin coordination described for GPCRs.

A key role of CCR9 is coordinating immune cell localization and positioning in the small intestine, which contributes to immune homeostasis and inflammatory responses. Receptor dysregulation is associated with a variety of inflammatory bowel diseases (IBDs), as well as cardiovascular disease, arthritis, and others [6–8]. Additionally, CCR9 and CCL25 overexpression has been observed in malignant tumors and is correlated with metastasis to the colon [9,10]. Therefore, CCR9 has emerged as a potential therapeutic target for various diseases [7,11,12]. Antagonizing CCR9 by small molecules effectively decreased inflammation in the colon in mouse models [13,14], however this effect was not replicated in clinical trials [15]. Pharmacological inhibition of the receptor by specific monoclonal antibodies effectively inhibited T lymphoblastic leukaemia tumor growth in pre-clinical studies [16,17]. Therefore, addressing CCR9-related pathophysiology may require more specific approaches and further elucidating native receptor regulation may provide insights into alternative strategies.

Here we present a detailed investigation of the signaling and regulation of CCR9 by the classical GPCR mediators, including G proteins, GRKs, and β-arrestins. Our results reveal CCR9 activation results in limited G protein signaling that is rapidly terminated. Rather than arrestins, GRK phosphorylation of specific C-terminal serine and threonine residues mediate the loss of G protein coupling. Phosphorylation appears to directly interfere with stable and productive G protein coupling. We propose that the activation of CCR9 is tightly regulated by GRK phosphorylation via an arrestin-independent mechanism.

## Results

### CCR9 robustly recruits mini-G_i_, but shows limited activation of the full heterotrimer

Activation of CCR9 drives immune cell migration through the activation of G_i_ and G_q_ signaling pathways [18–24]. G protein coupling can be generally described as several sequential and discrete steps, which we have simplified into three critical events. First the G protein engages the active receptor, then G protein activation leads to separation of the Gα from the Gβγ subunits, and finally the active G protein coordinates downstream signaling responses. Initial recruitment was tested by tracking bioluminescence resonance energy transfer (BRET) between CCR9 with a C-terminal luciferase (CCR9-RlucII) and a fluorophore fused to a mini-G (mG_x_) [25]. Mini-G proteins are engineered G protein mimetics containing the Ras domain of Gα_s_ and with the specific interacting residues of G_i_ or G_q_. When CCR9 is activated by CCL25, it leads to a rapid and robust recruitment of mG_i_ to the receptor with similar kinetics as observed for CXCR4, a well-studied and efficient activator of G_i_ proteins, stimulated by CXCL12 (Fig. 1A). In stark contrast, ACKR4, the other native receptor binding CCL25, shows no change in BRET between the receptor and mG_i_, consistent with the atypical nature of its activation and lack of G protein coupling [26]. Effective recruitment of mG_q_ is also observed for CCR9 (Fig. S1A).

Following recruitment, the G protein is activated which leads to dissociation of the Gα and Gβγ subunits. By introducing BRET sensors into the heterotrimer between the Gα and Gβγ subunits, G protein activation can be tracked by a decrease in signal following activation by a GPCR [27]. Despite showing robust mG_i_ recruitment (Fig. 1A), stimulation of CCR9 by CCL25 only produces a modest decrease in BRET between the G protein subunits (ΔBRET ∼2%) that rapidly returns to basal levels within 10 min (Fig. 1B). CXCR4, in contrast, shows a proportionally large signal (ΔBRET ∼10%) which is sustained for the duration of the experiment. A similarly muted response to CCR9 activation is observed with G_q_ activation (Fig. S1B) suggesting this weak activation, despite strong recruitment, is a general feature of CCR9 signaling and not specific to certain G proteins.

The limited G protein activation is further observed in downstream signaling by G_i_ proteins. G_i_ activation inhibits cAMP production. To characterize this response, the inhibition of forskolin-induced cAMP production by activated chemokine receptor was tracked using a BRET-based cAMP biosensor (CAMYEL) [28]. Like with the G protein dissociation, CCR9 activation by CCL25 had a minor effect on cAMP compared to mock transfected cells, while CXCR4 activated by CXCL12 substantially reduces forskolin-promoted cAMP production (Fig. 1C).

**Figure 1.**
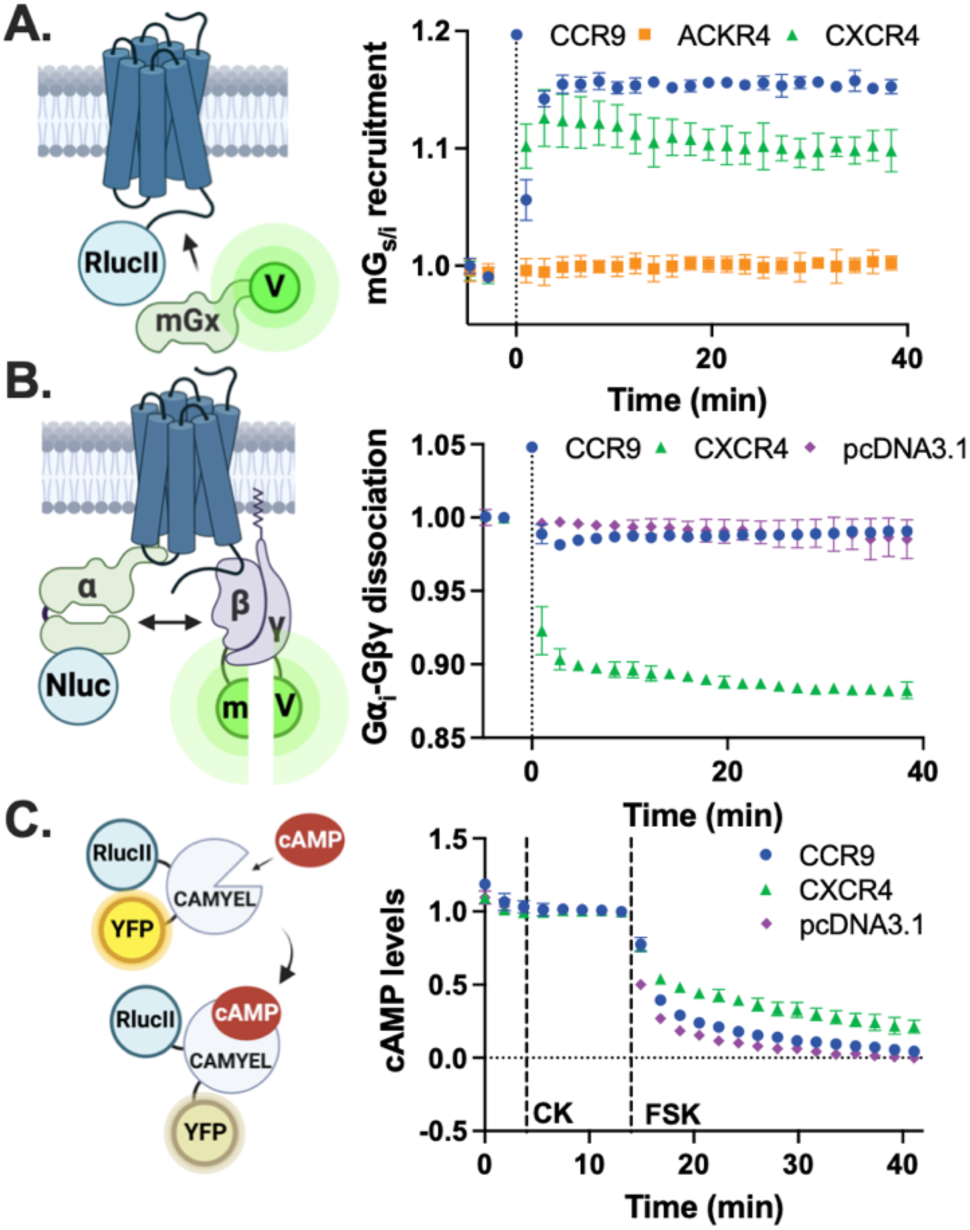
CCR9 recruits G_i_ proteins but shows limited heterotrimer activation. (A) Ligand-induced recruitment of mini-G⍺_i_ to CCR9-RlucII, ACKR4-RlucII, and CXCR4-RlucII in HEK293 cells measured by BRET following stimulation with 100 nM chemokine. (B) Ligand-induced activation of G_i_ proteins measured as dissociation of G⍺_i_-Nluc and Gβγ-split-mVenus (Gβγ-smV) in HEK293 cells by CCR9 and CXCR4 upon stimulation of 100 nM CCL25 or CXCL12, respectively. (C) Inhibition of cAMP production by CCR9 and CXCR4 in HEK293 using the BRET-based cAMP sensor CAMYEL. Cells were first treated with 100 nM chemokine (CK, CCL25 or CXCL12) and then cAMP production was stimulated by addition of 10 µM forskolin (FSK). Values represent mean ± SD of three independent experiments performed in triplicate.

### The CCR9 C-terminus suppresses CCL25-induced G protein coupling

The rapid CCR9 signal attenuation suggests that a fast inhibitory mechanism suppresses the coupling of G proteins to the active receptor. The GPCR C-terminus plays a critical role in canonical signal termination by coordinating arrestin interactions following GRK phosphorylation and has recently been described in some cases as autoinhibitory [29–31] and therefore may contribute to the poor G protein activation by CCR9. To resolve whether the C-terminus plays a role in regulation of CCR9 signaling, the receptor was truncated after position V334 (CCR9 ΔCT, Fig.2A). Consistent with an inhibitory function, CCR9 ΔCT showed robust and sustained G protein dissociation following CCL25 stimulation (Fig. 2B). The truncated receptor shows a greater maximal change in BRET (ΔBRET ∼7% v. ∼2%) and a 6-fold increase in total G protein activation as determined by the area over the dissociation curve (AOC) (Fig. 2C). Unlike other receptors [29,31,32], only the agonist-promoted G protein activation is enhanced for CCR9 ΔCT and constitutive activity is not altered (Fig. S2). Additionally, the surface expression of CCR9 ΔCT is only slightly lower than that of WT CCR9 (Fig. S3). These results suggest that the limited G protein activation by CCR9 is due to features of the receptor C-terminus suppressing signaling by the activated receptor.

**Figure 2.**
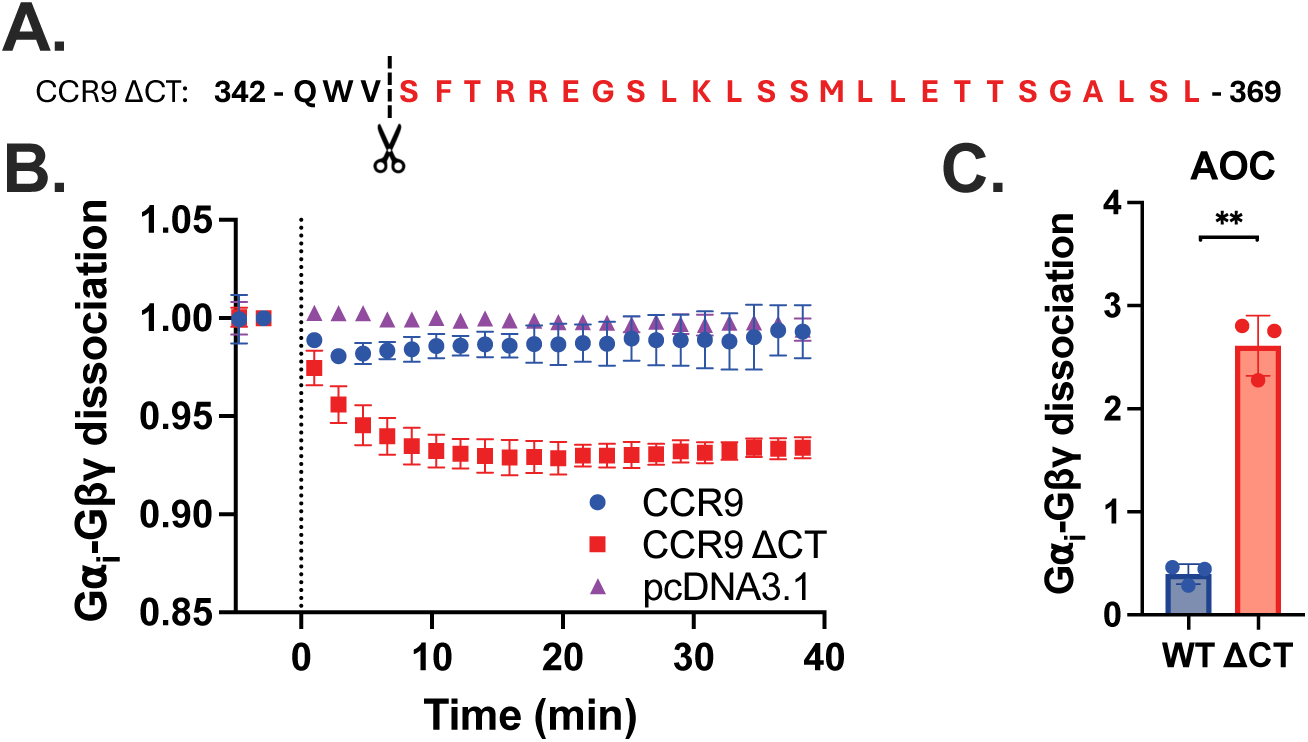
Truncating the CCR9 C-terminus significantly improves CCL25-induced G_i_ coupling. (A) Ligand-induced activation of G_i_ proteins measured as dissociation of G⍺_i_-Nluc and Gβγ-smV in HEK293 cells by CCR9 and CCR9 ΔCT upon stimulation of 100 nM CCL25. (B) Quantification of G⍺_i_-Gβγ dissociation by integration of the area over the BRET curves. Values represent mean ±SD of three independent experiments performed in triplicate, **P<0.001.

### Arrestins do not attenuate CCR9-mediated G protein signaling

GPCR C-termini canonically coordinate signal termination by acting as a scaffold for GRK phosphorylation, which in turn promotes arrestin recruitment to impede further G protein coupling [33,34]. As the CCR9 C-terminus appears to be critical for the rapid suppression of signaling, we evaluated how arrestins influence the coupling of G proteins to CCR9 by tracking the activation of G proteins through G protein dissociation in β-arrestin1/2 CRISPR knockout cells (Δβarr1/2) [35]. Prior to evaluating the impact of arrestins on G proteins, we first assessed β-arrestin2 recruitment to CCL25-stimulated CCR9 by BRET between CCR9-RlucII and GFP-tagged arrestin (GFP10-βarr2). Agonist addition leads to a rapid association of arrestin to CCR9 that peaks at ∼4 min and then slowly decreases to ∼50% the peak signal (Fig.3A). G_i_ protein dissociation by CCL25 stimulation of CCR9 in Δβarr1/2 cells shows identical levels of activation as in WT cells and do not match the enhanced activation with CCR9 ΔCT (Fig. 3B,C v 2B,C). Activation of G_q_ proteins were also unaffected by the absence of arrestins (Fig. S4). These data suggest that the C-terminal suppression of CCR9 signaling is not due to arrestin coordination.

**Figure 3.**
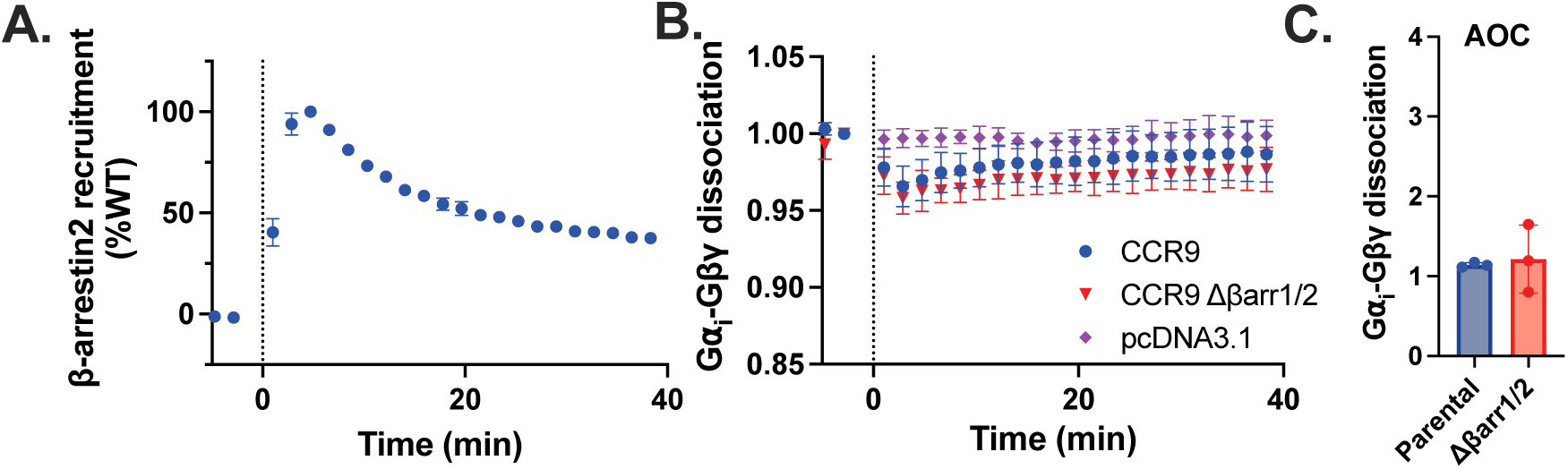
Arrestins do not impact CCR9 coupling to G proteins. (A) CCL25-induced recruitment of GFP10-βarr2 towards CCR9-RlucII detected by BRET over time following stimulation with 200 nM chemokine. (B) Ligand-induced activation of G_i_ proteins measured as dissociation of G⍺_i_-Nluc and Gβγ-smV in HEK293 Parental and HEK293 Δβarr1/2 cells by CCR9 upon stimulation with 100 nM CCL25. (C) Quantification of G⍺_i_-Gβγ dissociation in Parental and Δβarr1/2 cells by integration of the area over the BRET curves. Values represent mean ± SD of three independent experiments performed in triplicate.

### Phosphorylation of CCR9 by GRKs suppresses G protein activation

In addition to coordinating arrestin recruitment, the GPCR C-terminus undergoes phosphorylation at serine and threonine residues by GRKs, which drives internalization independent of arrestins in some cases [36–38]. Therefore, GRKs can influence GPCR activity in a direct manner and may coordinate the C-terminal suppression of CCR9. To determine the contributions of the four ubiquitously expressed GRKs (GRK2, 3, 5, and 6) to CCR9 phosphorylation, the recruitment of β-arrestin2 to CCR9 was measured in GRK family CRISPR knockout cells (ΔGRK2/3, ΔGRK5/6, and ΔGRK2/3/5/6) using a BRET recruitment assay (Fig. 4A,B) [39]. Arrestins require phosphorylation of the GPCR C-terminus for recruitment and these interactions are sensitive to specific phosphorylation patterns. Thus, changes to arrestin binding patterns correspond to changes in the receptor phosphorylation state and pattern. Arrestin recruitment is only partially impacted in ΔGRK2/3 or ΔGRK5/6 cells, suggesting little preference for CCR9 phosphorylation between the major GRK families. In ΔGRK2/3/5/6 cells, the recruitment is nearly eliminated, confirming that GRK phosphorylation is essential for CCR9-arrestin interactions. G protein activation by CCR9 in the ΔGRK2/3/5/6 cells, tracked by BRET through heterotrimer dissociation, revealed robust and sustained signaling (Fig. 4C) with a profile similar to CCR9 ΔCT (Fig. 2A). The ∼6-fold increase in the AOC by CCR9-mediated activation of G_i_ in the absence of GRKs matches the increase for the CCR9 ΔCT mutant (Fig.2B), suggesting that phosphorylation of the receptor C-terminus is a determining factor by which the domain drastically diminishes G protein coupling. A similar ∼6-fold increase is seen for G_q_-coupling towards CCR9 in ΔGRK2/3/5/6 cells (Fig. S5).

**Figure 4.**
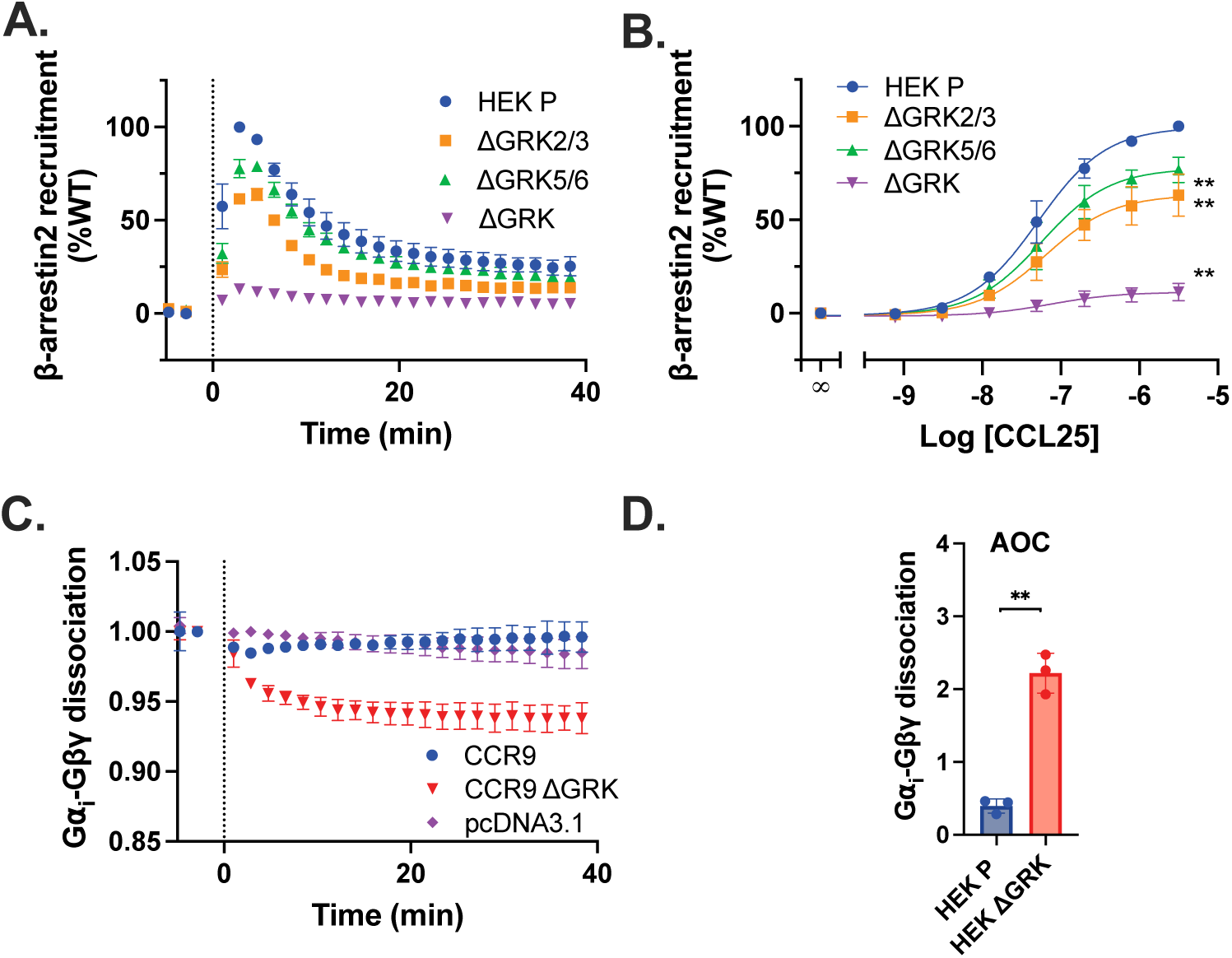
G protein activation by CCR9 is enhanced in the absence of GRKs. CCL25-induced recruitment of GFP10-βarr2 towards CCR9-RlucII detected by BRET over time following stimulation with 200 nM chemokine (A) or across a titration of CCL25 concentrations (B) in HEK293, ΔGRK2/3, ΔGRK5/6, and ΔGRK2/3/5/6 (ΔGRK) cells. (C) Ligand-induced activation of G_i_ proteins measured as dissociation of G⍺_i_-Nluc and Gβγ-smV in HEK293 ΔGRK cells by upon stimulation with 100 nM CCL25. WT data is repeated from figure 1B for comparison. (D) Quantification of G⍺_i_-Gβγ dissociation in Parental and ΔGRK cells by integration of the area over the BRET curves. Values represent mean ± SD of three independent experiments performed in triplicate. Statistical significance was determined by an unpaired t-test (AOC) or using the extra sum-of squares F test (DRC), **P<0.001.

### Phosphorylation of C-terminal sites proximal to CCR9 attenuates activation of G proteins

GRK phosphorylation of GPCR C-termini can produce specific modification patterns which are proposed to coordinate specific responses via arrestin [40]. The clear involvement of GRKs in suppressing CCR9 signaling presents the possibility that this regulation may be due to phosphorylation ‘barcodes’ on the receptor C-terminus. To resolve which sites are responsible for signal attenuation, two clusters of serines and threonines on the CCR9 C-terminus (C1, S335A/T347A/S352A/S356A/S357A and C2, T362A/T363A/S364A/S368A) were mutated to alanines to selectively prevent specific GRK phosphorylation (Fig. 5A). The loss of phosphorylation was validated by β-arrestin2 recruitment measured by BRET. Both CCR9 C1 and CCR9 C2 showed impaired β-arrestin2 recruitment, with the latter having a larger effect (Fig. 5B,C). Indicating that both the proximal and the terminal phosphorylation sites are involved in the recruitment of arrestins. As expected, mutation of all the phosphorylation sites on the C-tail (C1+C2) almost completely eliminated the CCL25-induced arrestin recruitment. Next, we tested G_i_ dissociation by the CCR9 ST/A mutants. Both CCR9 C1 and C1+C2 showed a ∼2-fold increase in G protein dissociation (Fig. 5D,E), while C2 did not alter the rapid suppression of CCR9 signaling. An identical pattern was observed for suppression of cAMP production, with both CCR9 C1 and C1+C2 showing decreased levels (Fig. 5F,G). No correlation was observed between direct GRK recruitment towards the receptor and G protein coupling (Fig. S6). Signal attenuation is regulated by the proximal (C1), but not terminal (C2), phosphorylation sites, while GRK recruitment were similar for both constructs. This suggests that the enhancement observed for C1 and C1+C2 is specifically due to changes in C-terminal phosphorylation and not recruitment of the GRKs. Mutations to CCR9 also had no effect on the basal G_i_ dissociation and cAMP levels (Fig. S7A,B). In GRK deficient cell lines we measured a ∼6-fold increase of G protein recruitment of CCR9, while eliminating the proximal phosphorylation sites only showed a ∼2-fold increase, suggesting inhibition by GRKs is not solely due to phosphorylation. This discrepancy could not be explained by altered surface expression of the CCR9 mutants, as differences in the presented receptor levels did not correlate with enhancements to G protein activation (Fig. S8). Additionally, attempts to increase the number of receptors on the plasma membrane by amplifying the amount of transfected DNA did not change the effect on G protein coupling (Fig. S9).

**Figure 5.**
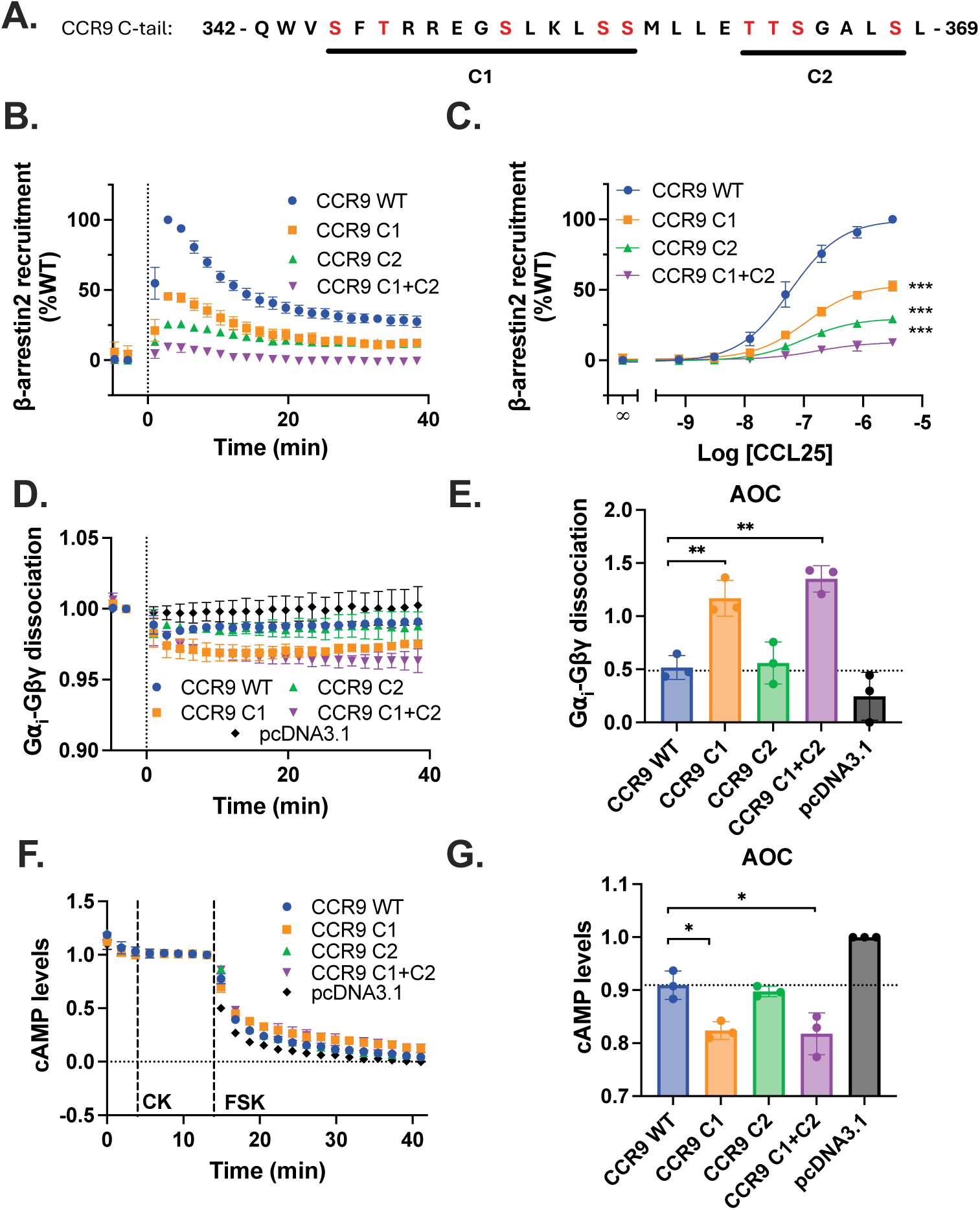
Proximal phosphorylation sites regulate G protein activation. (A) Schematical view of CCR9 C-tail mutants. Highlighted amino acids were mutated into alanines. CCL25-induced recruitment of GFP10-βarr2 towards CCR9-RlucII detected by BRET over time following stimulation with 200 nM chemokine (B) or across a titration of CCL25 concentrations (C) in HEK293 cells. (D) Ligand-induced activation of G_i_ proteins measured as dissociation of G⍺_i_-Nluc and Gβγ-smV in HEK293 cells after stimulation of CCR9 WT and ST/A mutants with 100 nM chemokine. WT data is repeated from figure 1B for comparison. (E) Quantification of G⍺_i_-Gβγ dissociation by integration of the area over the BRET curves. (F) Inhibition of cAMP production by CCR9 in HEK293 using the BRET-based cAMP sensor CAMYEL. Cells were first treated with 100 nM CCL25 (CK) and then cAMP production was induced by addition of 10 µM FSK. WT data is repeated from figure 1C for comparison. (G) Quantification of cAMP levels by integration of the area over the BRET curves. Values represent mean ± SD of three independent experiments performed in triplicate. Statistical significance was determined by one-way Browns-Forsythe & Welch ANOVA followed by an unpaired t-test (AOC) or using the extra sum-of squares F test (DRC). *P<0.05, **P<0.001, ***P<0.0001.

### CCR9 internalization does not attenuate G protein signal attenuation

GRK phosphorylation of GPCRs generally mediates receptor internalization through both arrestin-dependent and -independent mechanisms and is often considered a key step in receptor deactivation. Recent reports have shown that some receptors continue to signal from endosomal bodies [41], however it is not known if this is the case for CCR9. Therefore, if CCR9 lacks intracellular signaling capabilities, phosphorylation-mediated internalization may account for the observed signal termination. Translocation of the receptor after ligand stimulation was monitored by bystander BRET between CCR9-RlucII and an acceptor anchored at the plasma membrane by a CAAX motif (rGFP-CAAX) [42]. In WT cells, CCR9 shows robust and sustained translocation away from the plasma membrane with stimulation by CCL25 (Fig. 6A). In comparison, the level of CCR9 internalization in the Δβarr1/2 cells is significantly reduced to ∼50% that of WT cells. In contrast, CCL25-mediated CCR9 internalization is absent in ΔGRK2/3/5/6 cells, but nearly WT-like in knock-outs of GRK2/3 and GRK5/6 (Fig. 6B). These results would be consistent with CCR9 translocation being a driver of G protein signal attenuation and only receptors on the surface are capable of coupling. However, internalization of the phospho-substitution mutants reveals that trafficking of CCR9 is mainly mediated by the phosphorylation of the C2 sites (Fig. 6C). In the absence of these distal phosphorylation sites, CCR9 no longer shows agonist-mediated internalization. CCR9 C1 still internalizes with CCL25 stimulation albeit 50% that of WT. As loss of the C1 phosphorylation leads to enhanced CCR9 signaling and maintains partial internalization, the relocalization of CCR9 with CCL25 activation does not account for the acute signal suppression observed in Fig. 1.

**Figure 6.**
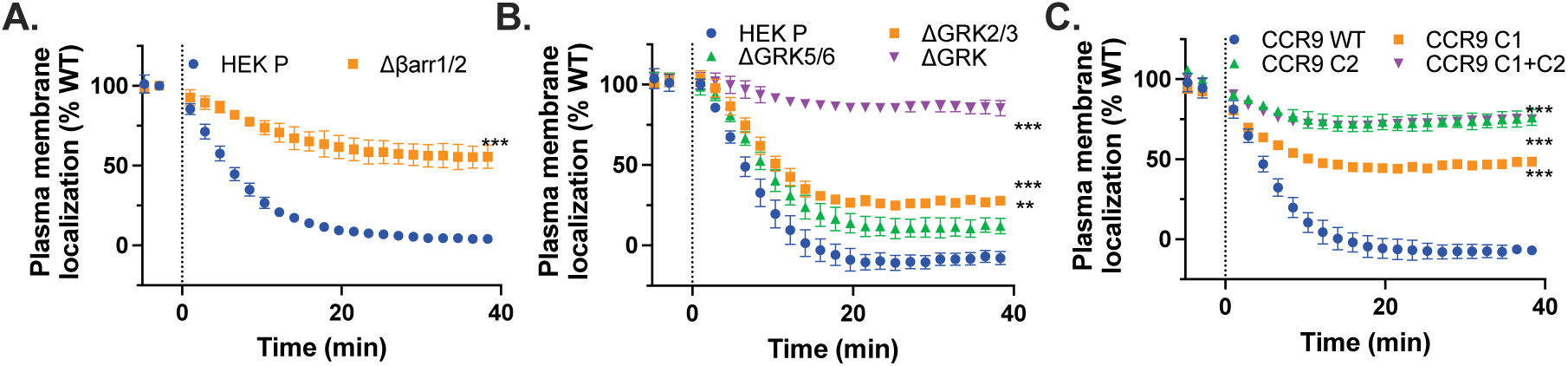
Internalization not the driving factor for desensitization. (A) CCL25-induced internalization measured as bystander BRET between CCR9-RlucII and rGFP-CAAX over time following stimulation with 200 nM chemokine in HEK293 Parental and Δβarr1/2 cells (A), in HEK293 Parental, ΔGRK2/3, ΔGRK5/6 and ΔGRK2/3/5/6 (ΔGRK) cells (B), and of the CCR9 C-tail mutants in HEK293 cells (C). Values represent mean ± SD of three independent experiments performed in triplicate. Statistical significance was determined by an unpaired t-test, **P<0.001, ***P<0.0001.

### CCR9 phosphorylation disrupts interactions with G proteins

Having eliminated the more common phosphorylation-mediated mechanisms for GPCR deactivation, we asked how does proximal phosphorylation inhibit G protein activation? One possible explanation is that the C-terminal phosphorylation sites are involved in stabilizing the CCR9:G protein ternary complex. Most structures of GPCRs and G proteins lack information on the receptor C-terminus with a few notable exceptions modeling the proximal receptor C-terminus in the cleft between the Gα and Gβ subunits [43]. To resolve how phosphorylation impacts the CCR9:G protein complex formation, the interaction between the receptor and the effector was tracked by BRET. First, the effect of CCL25-activation on CCR9-Gα_i_ interactions was monitored between a receptor with a C-terminal mVenus acceptor and a Nluc on Gα_i_ (Fig. 7A). Full length Gα_i_ was used for this assay instead of the mG_i_ from Fig. 1A to consider all interactions of the signaling heterotrimer. In WT cells, CCR9 shows rapid dissociation from G⍺_i_ with CCL25-stimulation (Fig. 7B,C). The rate of dissociation outpaces the rate of internalization, suggesting that it is unlikely that the effect is due to the receptor moving away from the heterotrimer (Fig. 7B v Fig. 6). This suggests that G⍺_i_ precomplexes with CCR9 and receptor activation decreases this association. The CCL25-promoted dissociation between Gα_i_ and CCR9 is significantly reduced in ΔGRK2/3/5/6 cells where agonist-mediated phosphorylation is no longer possible, indicating that phosphorylation partially disrupts the interaction with Gα_i_. The dissociative response of the full-length Gα_i_ with CCL25 stimulation conflicts with the association observed for mG_i_ (Fig. 1A). This is likely due to the engineered features of mG_i_, lack of lipidation or the Gα_s_ scaffold, limiting potential receptor interactions compared to native Gα_i_ [44].

GPCR:G protein structures containing receptor C-terminal density suggest that interactions are also made with the Gβ subunit of the heterotrimer [43]. The interaction between CCR9-RlucII and Gβγ-smV was tracked by BRET (Fig. 7D). Similar to Gα_i_, Gβγ quickly dissociated from CCR9 in WT cells following CCL25 stimulation (Fig. 7E,F) at a faster rate than receptor internalization (Fig. 6). Unlike the partial effect observed for Gα_i_ in ΔGRK2/3/5/6 cells, the Gβγ:CCR9 BRET signal does not decrease with agonist stimulation. This suggests that GRK phosphorylation of CCR9 leads to a destabilization of interactions with the G protein heterotrimer. Without phosphorylation, the Gβγ dimer remains associated with CCR9 during signal propagation, potentially acting to coordinate further G protein coupling.

**Figure 7.**
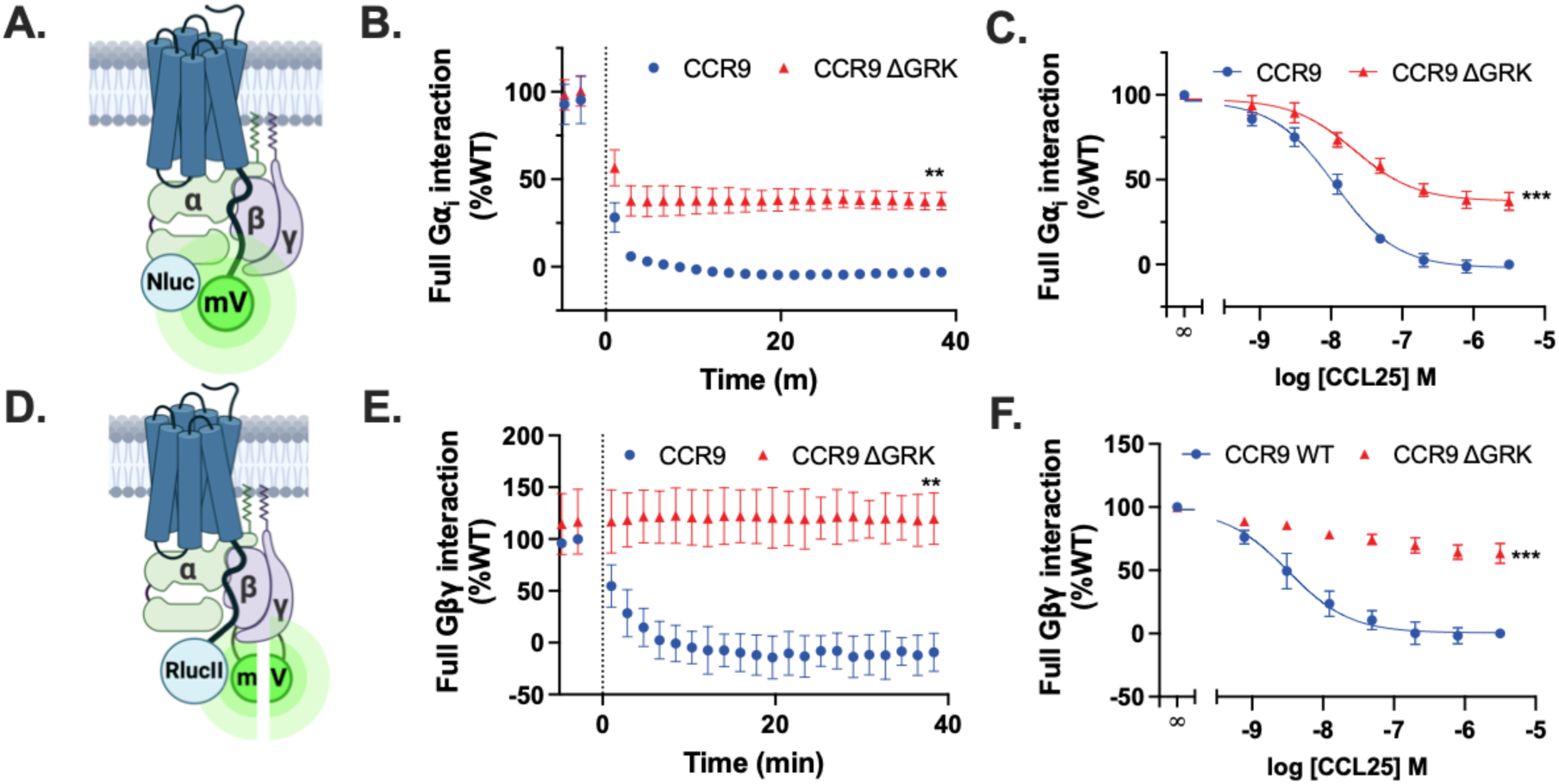
Proximal phosphorylation impairs C-tail interaction with G⍺/Gβγ. (A) Schematic illustration of BRET pairs CCR9-mVenus and G⍺_i_-Nluc. Ligand-induced interaction between G⍺_i_-Nluc and CCR9-mVenus in HEK293 and HEK293 ΔGRK2/3/5/6 (ΔGRK) cells measured by BRET over time following stimulation with 200 nM chemokine (B) or across a titration of CCL25 concentrations (C). (D) Schematic illustration of BRET pairs CCR9-RlucII and Gβγ-smV. Ligand-induced interaction between Gβγ-smV and CCR9-RlucII in HEK293 and HEK293 ΔGRK cells measured by BRET over time following stimulation with 200 nM chemokine (E) or across a titration of CCL25 concentrations (F). Values represent mean ± SD of three independent experiments performed in triplicate. Statistical significance was determined by an unpaired t-test (B, E) or using the extra sum-of squares F test (C, F). **P<0.001 ***P<0.0001.

**Figure 8.**
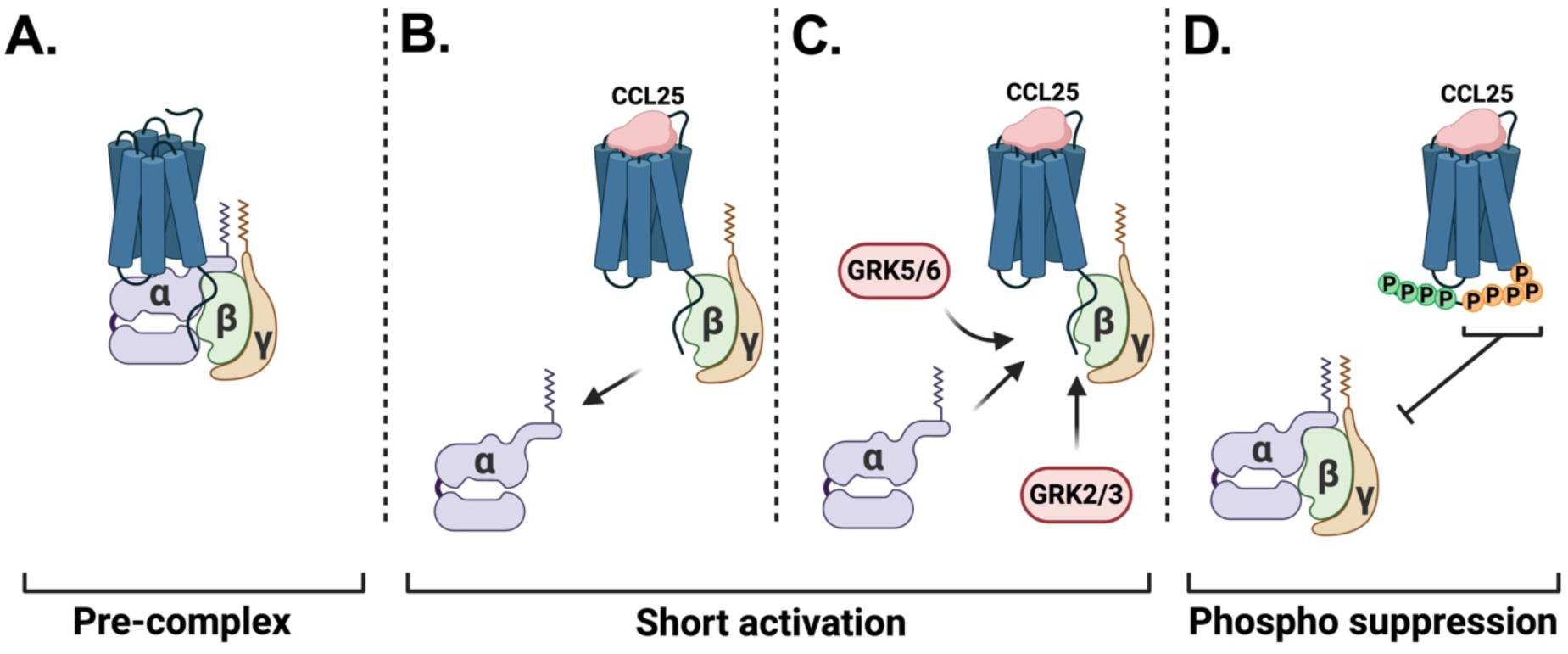
CCR9 pre-complexes with G proteins that is disturbed by proximal phosphorylation of the C-tail by GRKs upon receptor activation. (A) Empty CCR9 forms a pre-complex with the full G protein heterotrimer (Fig. 7). (B) Immediately upon chemokine binding, CCR9 is activated and initiates G protein signaling. The Gα unit dissociates to initiate downstream responses like cell migration, while the Gβγ remains associated with the activated receptor (Fig. 7B,E). (C) While activated, CCR9 is also primed for phosphorylation by GRKs. The kinases bind to the same interface as the G protein and compete for receptor interaction. (D) GRK phosphorylation of the proximal receptor C-terminus disrupts interactions and productive coupling of the G protein with CCR9. This leads to a dissociation of the CCR9-G protein complex (Fig. 7B,E), rapidly terminates G protein activation, and allows for the reassociation of the heterotrimer (Fig. 1B).

## Discussion

CCR9 mediates immune cell homeostasis in the gut and over or aberrant expression is tied to several inflammatory disorders and cancer metastasis. Therefore, suppressing receptor signaling is a promising therapeutical outcome, although direct antagonism of the receptor has proven difficult [15]. Here, we describe a non-canonical mechanism of CCR9 signal termination, whereby the classical model of arrestin inhibition of GPCRs is circumvented. Instead, GRK-mediated phosphorylation of the CCR9 C-terminus directly attenuates and suppresses G protein signaling. This allows for a limited signaling window while G proteins and GRKs compete for access to the activated receptor (Fig. 1B). Recruitment of GRKs leads to phosphorylation which inhibits further G protein signaling by interfering with productive coupling between the heterotrimer and the active GPCR (Fig. 7B,E) and leads to rapid loss of G protein coupling in our HEK293 system (Fig. 8). Specific phosphorylation sites mediate the observed attenuation, expanding the phosphorylation barcode paradigm beyond arrestins. Alternate systems of GPCR regulation may present new avenues for specific therapeutic intervention.

The phosphorylation barcode hypothesis suggests that specific patterns of phosphorylation influence the interactions with effectors into specific outcomes. Although this is often discussed from an unbiased perspective, the only effect generally considered is the impact of the phosphorylation barcode on interactions and activation states of arrestins. An implicit assumption of these analyses is often that arrestins are the sole translators of different phosphorylation patterns into cellular responses [45]. Here, we present evidence of arrestin-independent barcoding, where the phosphorylation alone is sufficient to terminate receptor signaling responses. Direct phospho-regulation of chemokine receptors is not a new concept. Efficient scavenging and internalization of the atypical chemokine receptor ACKR3 is wholly dependent on GRK phosphorylation, while arrestins are dispensable [37,46]. In the case of ACKR3, the mechanisms connecting phosphorylation to functional regulation are unknown and may involve previously undescribed interactions. With CCR9, the addition of phosphates appears to destabilize the GPCR:G protein complex, which may prevent sustained responses. Phosphorylation of the C-terminus may disrupt key interactions between the receptor and G protein through altering possible hydrogen bonding or changes to the electrostatics of the flexible domain. Importantly, this deactivation model does not require additional, unknown interactors and can be fully explained by proposed direct contacts between the receptor and G protein. Though other factors may contribute, we prefer to propose the simplest possible model where phosphorylation of specific serine and threonine residues within the C-terminus of CCR9 interfere with further G protein coupling thereby terminating G protein signaling.

Canonically, the C-terminus of GPCRs coordinates signal attenuation and termination by scaffolding arrestin to sterically impair G protein coupling. Recent studies have shown that the C-terminus of other GPCRs, including the prototypical GPCRs rhodopsin [47,48] and β_2_ adrenergic receptor [29], interacts with the cytoplasmic face of the receptor and interferes with G protein coupling. Truncation of the C-terminus leads to not only a large increase in stimulated G protein signaling, but also enhanced constitutive, unliganded G protein interactions. Structural and biophysical approaches have resolved interactions of the C-terminal tail with the receptor cytoplasmic face that is relieved with receptor activation [29,47]. In the case of CCR9, no change in constitutive activation is observed with ΔCT (Fig. S2), suggesting the CCR9 C-terminus lacks an autoinhibitory role. Curiously, the increase in signaling by other C-terminally truncated receptors was observed in the absence of arrestins [29]. This indicates a non-canonical arrestin-independent mechanism contributes to signal suppression in these other canonical GPCRs and may have been overlooked due to the dominance of the current paradigm, arrestin-mediated termination. In contrast, the rapid loss of CCR9 signaling is completely arrestin-independent and only driven by the receptor C-terminus and subsequent modifications. This suggests there may be degrees of relative contribution of arrestin-dependent and independent signal termination regulating GPCR signaling, with CCR9 being an extreme example of arrestin-independence.

Despite an abundance of structures of this complex being reported, only a handful resolve any C-terminal density. In these few structures, the C-terminus lays between the Gα and Gβ subunits of the G protein, forming contacts with both subunits [43,49]. In the structure of rhodopsin with its native G protein transducin, the residues that form these interactions include the phosphorylation sites S334 and T336 on the receptor which contact C271, D290, and D291 on Gβ. When these sites are mutated to alanines, the receptor shows greater and more sustained G protein coupling [50]. The most proximal phosphorylation sites on CCR9, S345 and T347, have identical positioning as S334 and T336 on rhodopsin relative to the conserved 8.50 position in helix 8 (H8) (Fig. S10). Phosphorylation adds a negative charge to these residues which may conflict with the electronegative aspartates forming the Gβ interface. Preventing phosphorylation leads to not only greater and sustained G protein activation, but also stabilizes the interactions between CCR9 and Gβγ (Fig. 7E). Increased stabilization of the CCR9:G protein complex may contribute to the enhanced G protein activation. In the case of rhodopsin, enhanced G protein signaling from ST/A receptor mutations was accredited to loss of arrestin interactions [51]. The results with CCR9 suggests there may also be a direct contribution of phosphorylation.

How does the muted CCR9 signaling translate to a physiological context? The HEK293 overexpression system used here is naturally far removed from the native immune cells environment. It is well established that CCR9 coupling to G proteins drives downstream signaling and cell migration, however, neither activation at the level of the G protein nor how the signal is terminated have been extensively investigated [22,24,52,53]. In our WT cell background, only limited signaling is observed, which is quickly suppressed with phosphorylation. However, this transient G protein activation may be sufficient for CCL25-driven cell migration and the rapid attenuation may have been evolved as a tighter regulation mechanism. One could draw comparisons to rhodopsin, where receptor activation and deactivation is tightly controlled to improve sensitivity [54]. By limiting the magnitude and length of signaling, CCR9 could provide specific and detailed positioning information for cells in the thymus and gut. Alternatively, the native system of CCR9 may have different GRK availability for phosphorylation. GRK levels in immune cells have been shown to be mediated by cytokine treatment, which in turn alters the localization of CCR9 proteins [55]. Thus, allowing for dynamic control and regulation over CCR9 signaling.

In conclusion, CCR9 activation is followed by rapid signal attenuation due to specific phosphorylation on the receptor C-terminus. In contrast to canonical GPCR mechanisms, the phospho-regulation of CCR9 is not coordinated by arrestins sterically blocking G protein coupling. Instead, phosphorylation directly destabilizes productive G protein coupling. This non-canonical mechanism for signal termination may complicate efforts to drug the receptor. Other GPCRs likely display similar mechanisms, but are often overlooked or missed due to dominant arrestin contributions. Thus, deactivation of GPCRs is perhaps more complicated than the canonical picture and may confound efforts to antagonize these receptors.

## Experimental procedures

### Materials

Unless stated otherwise, all chemicals and reagents were purchased from Melford or Sigma-Aldrich. HEK293 Δβ-arrestin1/2 cells (HEK293 Δβarr1/2) and corresponding parental cell lines were kindly gifted from Asuka Inoue (Tohoku University) [35]. HEK293 ΔGRK2/3, ΔGRK5/6, ΔGRK2/3/5/6, and corresponding parental cell lines were kindly gifted from Carsten Hoffmann (Friedrich-Schiller-Universität Jena) [56].

### DNA constructs and site-directed mutagenesis

Human CCR9 (1-369), ACKR4 (1-350), and CXCR4 (1-352) were cloned into pcDNA3.1 expression vector either alone or followed by a C-terminal *Renilla* luciferase II (RlucII), mVenus, or NanoLuc (Nluc). HA-H1R-Rluc8 [57], β-arrestin2-GFP10 (a gift from N. Heveker, Université de Montréal) [37], G⍺_i,3_ [58], G⍺_i_-Nluc, G⍺_q_-Nluc, GRK2-Nluc, GRK3-Nluc, GRK5-Nluc, GRK6-Nluc [26] (gifts from D. Legler, Biotechnology Institute Thurgau), split-mVenus(156-239)-Gβ_1_ [59], split-mVenus(1-155)-Gγ_2_ (gifts from N. Lambert, Augusta Univ.) [59], NES-Venus-mGs/i143 (mG_i_) [25,60], NES-Venus-mGq [25], mCitrine-EPAC-Rluc (CAMYEL) [28,58] and rGFP-CAAX (a gift from M. Bouvier, Université de Montréal) [42] were described previously. Site-directed mutagenesis was performed using Q5 Site-Directed Mutagenesis Kit (New England Biolabs) and confirmed by Sanger Sequencing.

### Chemokine purification from E. coli

The chemokines CCL25 and CXCL12 were expressed and purified from *E. Coli* as previously described [37,61]. Briefly, chemokine sequences were cloned into a pET21 vector together with N-terminal 8His-tag and enterokinase cleavage site. The generated cDNA was transformed into BL21(DE3)pLysS cells and expression was controlled by Isopropyl-β-D-thiogalactopyranoside (IPTG) induction. Inclusion bodies containing the chemokine were collected by sonication and dissolved in 50 mM Tris, 6 M guanidine-HCL, 50 mM NaCl, pH 8.0. The chemokine was purified using a Ni-NTA column, washed with 50 mM MES, 6 M guanidine-HCL, 50 mM NaCl, pH 6.0 and eluted with 50 mM acetate, 6 M guanidine-HCL, 50 mM NaCl, pH 4.0. Purified chemokine was denatured with 4 mM DTT and later refolded in 50 mM Tris, 500 mM arginine-HCl, 1 mM EDTA, 1 mM GSSG, pH 7.5. Refolded chemokine was dialysed in 20 mM Tris pH 8.0, 150 mM NaCl. Cleavage of the 8His-tag was done by addition of Enterokinase (New England Biolabs) and confirmed by SDS-PAGE and LC-MS. The cleaved material was polished on a Ni-NTA column, washed with 50 mM Tris pH 8.0 and eluted in 6 M guanidine, 50 mM MES, pH 6.0 and 6 M guanidine, 50 mM acetate, pH 4.0 as two separate fractions. The material was then bound to a C18 HPLC column (Gemini C18 110A, Phenomenex) (buffer A: 0.1% TFA, buffer B: Acetonitrile + 0.1% TFA) and eluted by a linear gradient of buffer B (5-95%). The chemokines were collected, lyophilized, and stored at -80 °C.

### Transfection HEK293 cells in suspension

Cells were cultured in Dulbecco’s Modified Eagle’s Medium (DMEM, Thermo Fisher Scientific) supplemented with 1% penicillin and streptomycin (Pen/Strep, Gibco) and 10% Fetal Bovine Serum (FBS, Bodinco) and maintained at 5% CO_2_ and 37 °C in a humidified atmosphere. HEK293 cells were transfected with a total of 2 µg DNA per 1 x 10^6^ cells using 6 µg polyethyleneimine (PEI, Polysciences Inc.) as a transfection reagent in 150 mM NaCl. The DNA-PEI mixture was mixed and incubated for 15 min at RT. HEK293 cells were detached using Trypsin/EDTA solution (Thermo Fisher Scientific) and resuspended in DMEM. Cells were counted and subsequently added to the DNA-PEI mixture. Cells were seeded at 30k/well in a white 96-well plate (Greiner) and incubated for 48 h unless stated otherwise.

### Mini-G⍺ recruitment by BRET

HEK293 cells were transfected with 50 ng CCR9-RlucII, ACKR4-RlucII, HA-CXCR4-RlucII or HA-H1R-Rluc8 and 250 ng NES-Venus-mG_s/i_143 or NES-Venus-mG_q_ up to an amount of 2 µg with pcDNA3.1 as described above. After 48 h incubation, cells were washed once with PBS and maintained in Hank’s Balanced Salt Solution (HBSS) supplemented with 0.1% Bovine Serum Albumin (BSA Fraction V, PanReac AppliChem). Cells were incubated with 5 µM of Coelenterazine-h (CTZ-h, Promega) for 5 min before 2 pre-read measurements were taken. Next, the cells were stimulated with 100 nM CCL25, CXCL12, or 10 µM Histamine. Bioluminescence was measured using 475-30 nm and 535-30 nm filters on a PHERAstar plate reader for 1h at 37 °C. BRET values were determined as the ratio of the red and blue luminescence readings (535-30/475-30). Results from three independent experiments were normalized to mock condition (no chemokine).

### G⍺-Gβγ dissociation by BRET

HEK293, HEK293 Δβarr1/2, HEK293 ΔGRK2/3/5/6 cells were transfected with 100 ng untagged receptor, 100 ng G⍺_i_-Nluc or G⍺_q_-Nluc, 500 ng Gβ-smV, and 500 ng Gγ-smV up to an amount of 2 µg with pcDNA3.1 as previously described. After 48 h incubation, cells were washed once with PBS and maintained in HBSS supplemented with 0.1% BSA. Cells were incubated with 2.5 µM of Furimazine (Promega) for 5 min before 2 pre-read measurements were taken. Stimulations, measurements and data analysis was performed as described previously (see Mini-G⍺ recruitment by BRET). Results from three independent experiments were normalized to mock condition (no chemokine). Quantification is done by integration of the area over the BRET curves (GraphPad Prism).

### cAMP measurements by BRET

HEK293 cells were transfected with 400 ng untagged receptor, 800 ng mCitrine-EPAC-Rluc and 400 ng G⍺_i,3_ up to an amount of 2 µg with pcDNA3.1 as described above. After 48 h incubation, cells were washed once with PBS and maintained in HBSS supplemented with 0.1% BSA. Cells were incubated with 5 µM of CTZ-h for 5 min before 3 pre-read measurements were taken. Next, the cells were stimulated with 100 nM CCL25 or CXCL12 and measured for 10 min. After incubation, cAMP levels were amplified with 10 µM Forskolin (FSK, Sigma-Aldrich) and bioluminescence was measured as described before (see Mini-G⍺ recruitment by BRET). Results from three independent experiments were normalized to pre-read to correct for basal differences.

### β-arrestin2 recruitment by BRET

HEK293, HEK293 ΔGRK2/3, HEK293 ΔGRK5/6, HEK293 ΔGRK2/3/5/6 cells were transfected with 50 ng CCR9-RlucII and 1 µg GFP10-β-arrestin2 up to an amount of 2 µg with pcDNA3.1 as described above. After 48 h incubation, cells were washed once with PBS and maintained in HBSS supplemented with 0.1% BSA. Cells were incubated with 5 µM of Prolume Purple (Prolume Ltd.) for 5 min before 3 pre-read measurements were taken. Next, the cells were stimulated with increasing concentrations of CCL25. Bioluminescence was measured at 410-80 nm and 515-30 nm using a PHERAstar plate reader for 1 h at 37 °C. BRET values were determined as the ratio of the red and blue luminescence readings (515-30/410-80). Results from three independent experiments were normalized to mock (no chemokine) or WT response as indicated.

### Active internalization assay by BRET

HEK293, HEK293 Δβarr1/2, HEK293 ΔGRK2/3, HEK293 ΔGRK5/6, HEK293 ΔGRK2/3/5/6 cells were transfected with 50 ng CCR9-RlucII and 200 ng rGFP-CAAX up to an amount of 2 µg with pcDNA3.1 as described above. Stimulations, measurements, and data analysis was performed as described above (see β-arrestin2 recruitment by BRET). Results from three independent experiments were normalized WT response as indicated and ΔBRET was calculated as BRET(CCL25)-BRET(mock).

### Full G⍺_i_ and Gβγ interaction by BRET

HEK293 or HEK293 ΔGRK2/3/5/6 cells were transfected with 250 ng CCR9-mVenus and 50 ng Gai-Nluc or 50 ng CCR9-RlucII and 250 ng Gβ-smV, and 250 ng Gγ-smV up to an amount of 2 µg with pcDNA3.1 as described above. After 48 h incubation, cells were washed once with PBS and maintained in HBSS supplemented with 0.1% BSA. Cells were incubated with 2.5 µM of Furimazine (G⍺_i_ assay) or 5 µM CTZ-h (Gβγ assay) for 5 min before 2 pre-read measurements were taken. Next, the cells were stimulated with increasing concentrations of CCL25. Measurements and data analysis was performed as described above (see Mini-G⍺ recruitment by BRET).

### Direct GRK recruitment by BRET

HEK293 cells were transfected with 50 ng GRK-Nluc and 250 ng CCR9-mVenus up to an amount of 2 µg with pcDNA3.1 as described above. Stimulations, measurements, and data analysis was performed as described above (see G⍺-Gβγ dissociation by BRET).

### Surface expression by flow cytometry

HEK293 cells were transfected with 100 ng CCR9 and up to an amount of 2 µg with pcDNA3.1 as previously described. After 48 h incubation, cells were washed once with PBS and transferred to a transparent 96-well plate (Guava compatible 96-well plate, Greiner) using Accutase (Thermo Fisher Scientific). From here on the cells were kept on ice constantly to prevent basal internalization and centrifuged at 350 x g for 3 min between each step. Cells were washed twice with cold FACS buffer (filtered PBS + 0.5% BSA). Next, cells were incubated with anti-hCCR9A antibody (R&D systems, MAB1791) for 1 h at 4 °C. After incubation, cells were washed twice with cold FACS buffer and consequently incubated with anti-mouse F(ab)2 IgG (H+L) PE-conjugated antibody (R&D systems, F0102B) for 1h in the dark at 4 °C. Cells were washed three times with cold FACS buffer and finally resuspended in FACS buffer. Mean yellow fluorescence was measured on the Guava easyCyte^TM^ (Cytek) and normalized to WT and pcDNA3.1 values.

### Statistical analysis

Statistical analyses were performed using GraphPad Prism 10. Bar and symbol representation, along with error bars, are described in figure legends. Scatter plots show the mean of three independent experiments, each measured in triplicate. For bar charts, the bars represent the mean of three independent experiments, whereas the points depict the mean values of the individual experiments measured in triplicate. All errors are reported as standard deviation (SD). The DRCs were plotted using a sigmoidal dose response model (log[agonist] vs. response, three parameters) using GraphPad Prism 10.

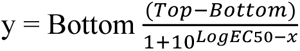

The Area Over the Curve (AOC) was determined in GraphPad Prism 10 (baseline = 1.0). Statistical significance of DRCs was determined using the extra sum-of squares F test. One-way Browns-Forsythe & Welch ANOVA followed by an unpaired t-test was done to determine statistical significance for multiple comparisons. All other statistical comparisons were done by an unpaired t-test.

## Data availability

All data for these studies are contained within this article.

## Supporting information

This article contains supporting information.

## Acknowledgements

We thank A. Inoue (Tohoku Univ.), C. Hoffmann (Friedrich-Schiller-Universität Jena), D. Legler (Biotechnology Institute Thurgau), and N. Lambert (Augusta Univ.) for the BRET constructs and cell lines used in our studies.

## Author contribution

T.D.L. and C.T.S. conceptualization; T.D.L. and C.T.S. formal analysis; M.J.S and C.T.S. funding acquisition; T.D.L. investigation; T.D.L and C.T.S. methodology; C.T.S. project administration; M.J.S. and C.T.S. resources; C.T.S. supervision; T.D.L and C.T.S. validation; T.D.L and C.T.S. visualization; T.D.L and C.T.S. writing – original draft; T.D.L., M.J.S., and C.T.S. writing – review & editing.

## Funding and additional information

This work was supported by the European Innovation Council through its Horizon Europe Pathfinder Open programme (Grant Agreement No. 101131014) and the Swiss State Secretariat for Education, Research and Innovation (SERI) [M.J.S., C.T.S.], the ONCORNET 2.0 (ONCOgenic Receptor Network of Excellence and Training 2.0) PhD training programme funded by the European Commission for a Marie Sklodowska Curie Actions (H2020-MSCA grant agreement 860229) [M.J.S].

