## Supplemental Information for "CCR9 signal termination is governed by an arrestin-independent phosphorylation mechanism"

**Supporting information**

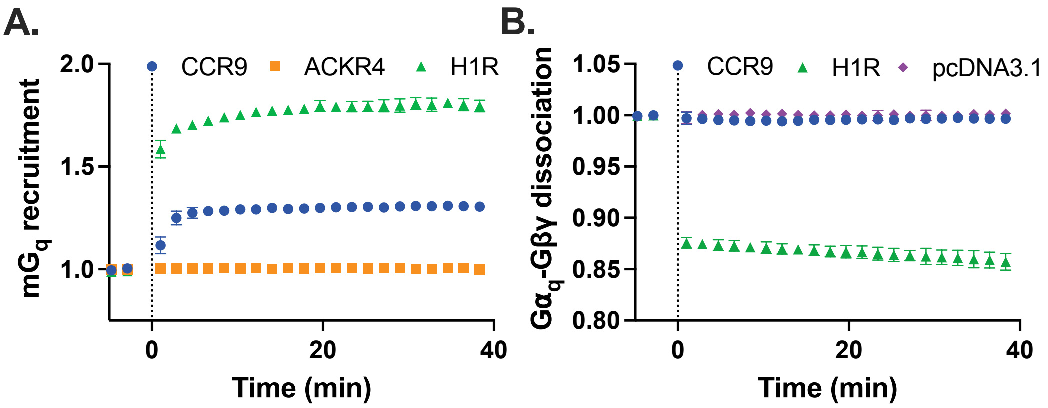


**Figure S1. CCR9 recruits $mG_q$ proteins but only weakly activates the $G_q$ heterotrimer.** (A) Ligand-induced recruitment of NES-Venus-$mG\alpha_q$ towards CCR9-RlucII, ACKR4-RlucII and H1R-Rluc8 in HEK293 cells detected by BRET over time following stimulation with 100 nM CCL25 or 10 µM Histamine. (B) Ligand-induced activation of $G_q$ proteins measured as dissociation of $G\alpha_q$-Nluc and Gβγ-smV in HEK293 cells in the presence of CCR9, H1R, and pcDNA3.1 upon stimulation of 100 nM CCL25 or 10 µM histamine. Values represent mean ± SD of three independent experiments performed in triplicate.

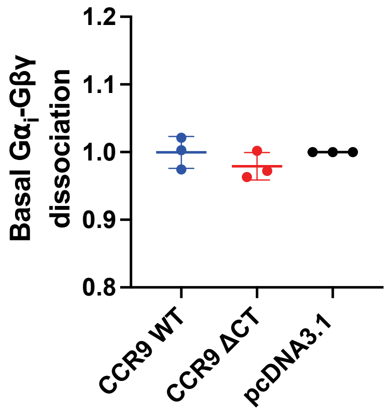


**Figure S2. CCR9 C-terminus does not inhibit constitutive G protein signaling.** Basal $G_i$ activation measured as dissociation of G$\alpha_i$-Nluc and Gβγ-smV in cells in the presence of CCR9, CCR9 ΔCT or empty pcDNA3.1.

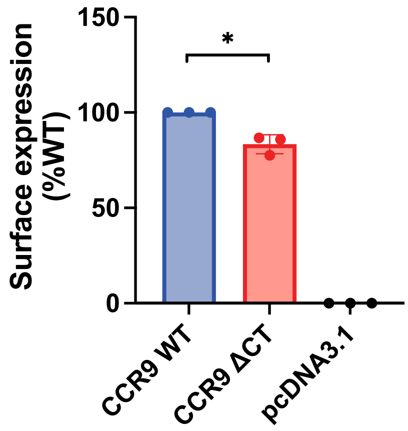


**Figure S3. Surface expression of CCR9 ΔCT is only slightly less than CCR9 WT.** Surface expression of untagged CCR9 in transfected HEK293 cells measured by flow cytometry. Bars represent mean ± SD of three independent experiments performed in triplicate. The average results of individual experiments are presented as points. Statistical significance was determined by a t-test. *P<0.05.

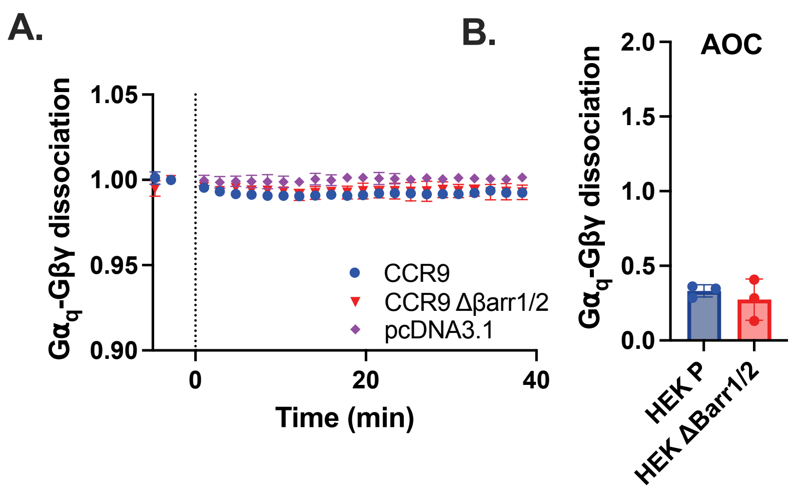


**Figure S4. Arrestins do not regulate suppression of CCR9 activation of $G_q$ proteins.** (A) Ligand-induced activation of $G_q$ proteins measured by the dissociation of $G\alpha_q$-Nluc and Gβγ-smV in HEK293 Δβarr1/2 cells in the presence of CCR9 or pcDNA3.1 upon stimulation of 100 nM CCL25 detected by BRET over time. (B) Quantification of $G\alpha_q$-Gβγ dissociation in Parental and Δβarr1/2 cells by integration of the area over the BRET curves. Values represent mean ± SD of three independent experiments performed in triplicate. Points represent the average of individual experiments.

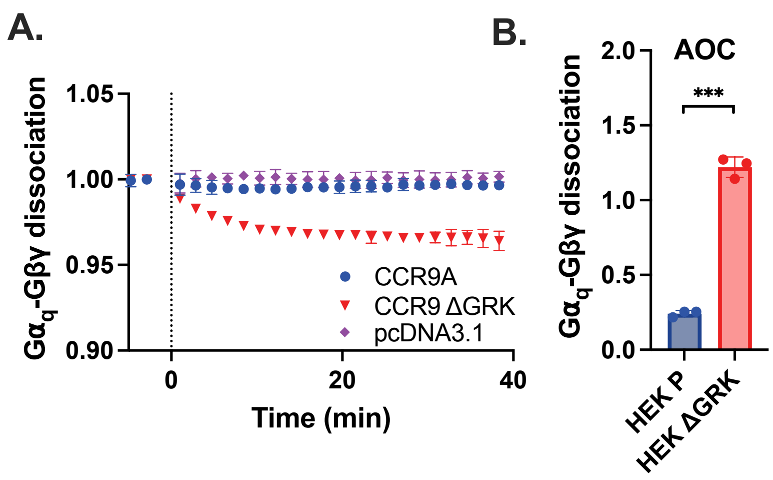


**Figure S5. CCR9 shows robust $G_q$ activation in GRK-deficient cells.** (A) Ligand-induced activation of $G_q$ proteins measured as dissociation of $G\alpha_q$-Nluc and Gβγ-smV in HEK293 ΔGRK2/3/5/6 (ΔGRK) cells in the presence of CCR9 or pcDNA3.1 upon stimulation of 100 nM CCL25 detected by BRET over time. (B) Quantification of $G\alpha_q$-Gβγ dissociation in Parental and ΔGRK cells by integration of the area over the BRET curves. Values represent mean ± SD of three independent experiments performed in triplicate. Points represent the average value from individual experiments. Statistical significance was determined by a t-test, ***P<0.0001.

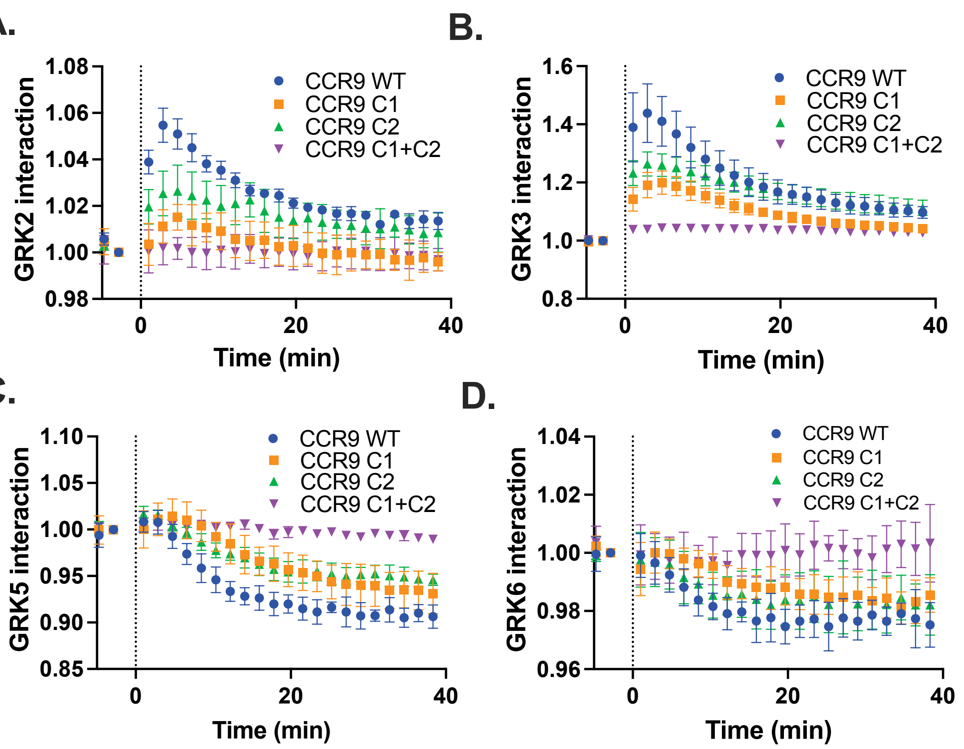


**Figure S6. Enhanced G protein coupling by CCR9 ST/A constructs does not correlate with differences in GRK recruitment.** Direct GRK recruitment of GRK2-Nluc (A), GRK3-Nluc (B), GRK5-Nluc (C), and GRK6-Nluc (D) towards CCR9-mVenus in HEK293 cells upon 200 nM CCL25 detected by BRET over time. Values represent mean ± SD of three independent experiments performed in triplicate.

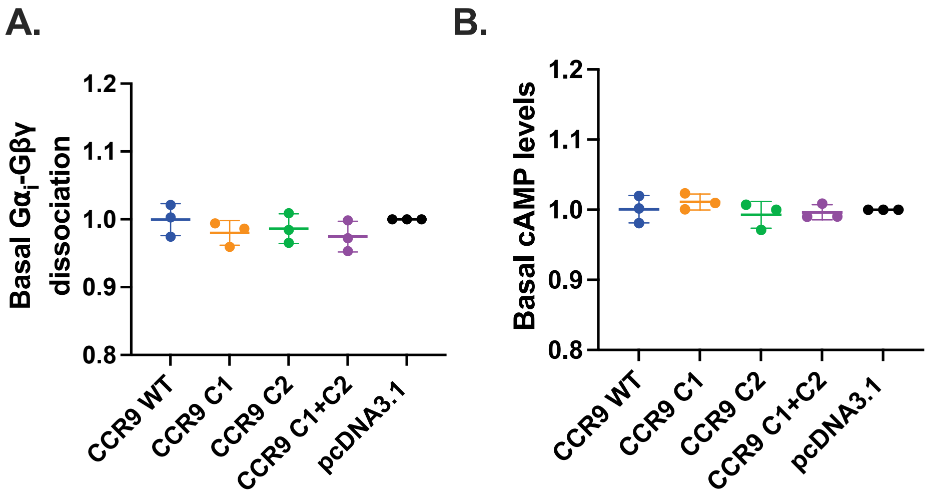


**Figure S7. ST/A CCR9 constructs do not constitutively activate G proteins.** (A) Basal $G_i$ activation measured as dissociation of $G\alpha_i$-Nluc and Gβγ-smV in cells in the presence of CCR9 or empty pcDNA3.1. CCR9 WT data is repeated from figure S2 for comparison. (B) Basal suppression of cAMP production by CCR9 or pcDNA3.1 in HEK293 using the BRET-based cAMP sensor CAMYEL. Values represent mean ± SD of three independent experiments performed in triplicate and normalized to pcDNA3.1 condition, while points represent the averages from the individual experiments.

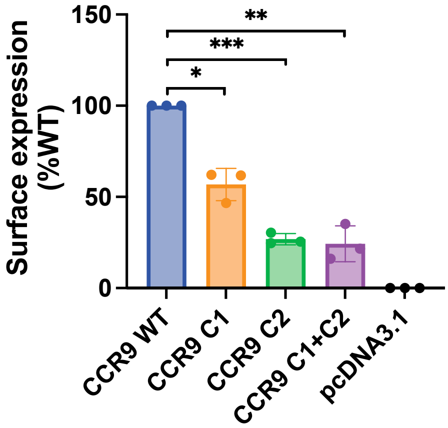


**Figure S8. Surface expression of CCR9 WT and ST/A mutants determined by flow cytometry.** Surface expression of untagged CCR9, CCR9 C1, CCR9 C2, and CCR9 C1+C2 in transfected HEK293 cells determined by flow cytometry. Values represent mean ± SD of three independent experiments performed in triplicate. Points present the average from individual experiments. CCR9 WT data is repeated from figure S3 for comparison. Statistical significance was determined by one-way Browns-Forsythe & Welch ANOVA followed by an unpaired t-test. *P<0.05, **P<0.001, ***P<0.0001

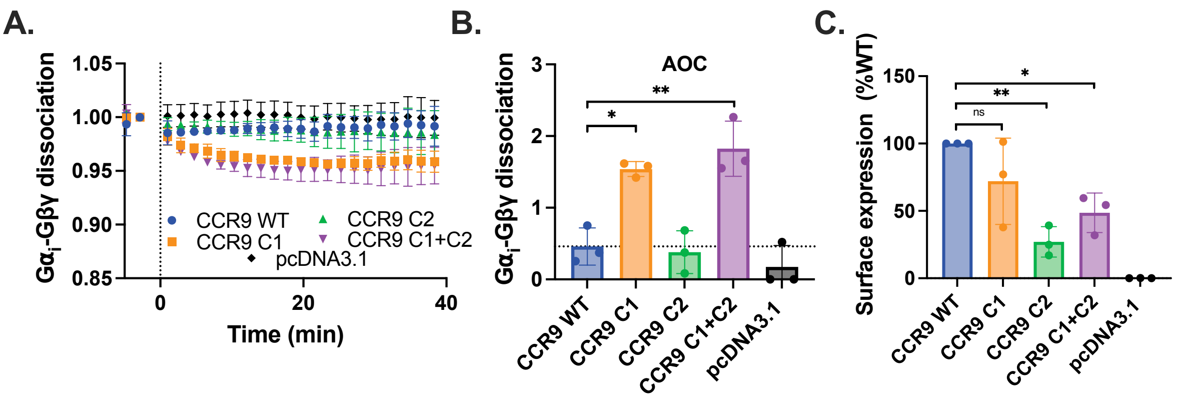


**Figure S9. Attempts to increase surface expression of CCR9 mutants did not change effect on G protein coupling.** To address the discrepancy in surface expression observed in Fig. S8, transfected DNA amounts were adjusted to attempt to match the surface expression of CCR9 WT (2xC1, 4xC2, 4xC1+C2). (A) Ligand-induced activation of $G_i$ proteins measured as dissociation of $G\alpha_i$-Nluc and Gβγ-smV in HEK293 cells in the presence of CCR9 or empty pcDNA3.1 upon stimulation of 100 nM chemokine. (B) Quantification of $G\alpha_i$-Gβγ dissociation by integration of the area over the BRET curves. (C) Surface expression of untagged CCR9, CCR9 C1, CCR9 C2, CCR9 C1+C2 in transfected HEK293 cells quantified by flow cytometry. Values represent mean ± SD of three independent experiments performed in triplicate. Points represent the average value from individual experiments. Statistical significance was determined by one-way Browns-Forsythe & Welch ANOVA followed by an unpaired t-test. *P<0.05, **P<0.001.

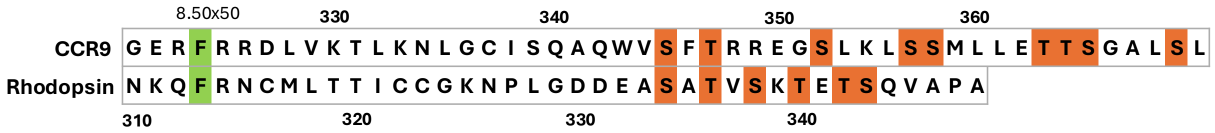


**Figure S10. Sequence alignment of CCR9 and rhodopsin.** C-terminal sequences were aligned by the conserved phenylalanine 8.50 in H8 of CCR9 and rhodopsin. Putative phosphorylation sites shown in orange.

**Table S11 Statistics**

| Fig. | Condition | Emax | | Log EC50 | | Statistical test |
|---|---|---|---|---|---|---|
| | | P-value | Value ± SD | P-value | Value ± SD | |
| **2b** | CCR9 WT | N.A | 0.40 ± 0.098 | N.A. | | t-test |
| | CCR9 ΔCT | 0.0028 | 2.61 ± 0.29 | | | |
| **4b** | HEK P | N.A. | 99.60 ± 3.89 | N.A | -7.29± 0.095 | Extra sum-of-squares F test |
| | ΔGRK2/3 | <0.0001 | 63.20 ± 11.24 | 0.39 | -7.20 ± 0.23 | |
| | ΔGRK5/6 | <0.0001 | 76.70 ± 6.82 | 0.62 | -7.25 ± 0.16 | |
| | ΔGRK | <0.0001 | 11.32 ± 4.61 | 0.59 | -7.13 ± 0.47 | |
| **4d** | HEK P | N.A | 0.40 ± 0.098 | N.A | | t-test |
| | HEK ΔGRK | 0.0036 | 2.22 ± 0.27 | | | |
| **5c** | CCR9 WT | N.A | 99.74 ± 4.00 | N.A | -7.23 ± 0.09 | Extra sum-of-squares F test |
| | CCR9 C1 | <0.0001 | 53.60 ± 2.79 | 0.0023 | -6.96 ± 0.11 | |
| | CCR9 C2 | <0.0001 | 29.85 ± 1.30 | 0.064 | -7,0 ± 0.095 | |
| | CCR9 C1+C2 | 0.0006 | 12.88 ± 2.62 | 0.1697 | -6.8 ± 0.45 | |
| **5e** | CCR9 WT | N.A | 0.52 ± 0.11 | N.A | | Browns-Forsythe & Welch ANOVA |
| | CCR9 C1 | 0.0077 | 1.17 ± 0.17 | | | |
| | CCR9 C2 | 0.77 | 0.56 ± 0.2 | | | |
| | CCR9 C1+C2 | 0.0011 | 1.35 ± 0.12 | | | |
| **5f** | CCR9 WT | N.A | 0,91 ± 0,03 | N.A | | Browns-Forsythe & Welch ANOVA |
| | CCR9 C1 | 0.014 | 0,82 ± 0,02 | | | |
| | CCR9 C2 | 0.53 | 0,90 ± 0,01 | | | |
| | CCR9 C1+C2 | 0.035 | 0,82 ± 0,04 | | | |
| **6a** | HEK P | N.A | N.A | N.A | | t-test |

| | | | | | | |
|---|---|---|---|---|---|---|
| | Δβarr1/2 | <0.0001 | 55.53 ± 7.33 | | | |
| **6b** | WT | N.A | N.A | N.A | | t-test |
| | ΔGRK2/3 | <0.0001 | 27.83 ± 2.56 | | | |
| | ΔGRK5/6 | 0.0052 | 12.00 ± 4.87 | | | |
| | ΔGRK | <0.0001 | 85.36 ± 4.76 | | | |
| **6c** | CCR9 WT | N.A | N.A | N.A | | t-test |
| | CCR9 C1 | <0.0001 | 48.48 ± 1.20 | | | |
| | CCR9 C2 | <0.0001 | 76.12 ± 4.87 | | | |
| | CCR9 C1+C2 | <0.0001 | 75.09 ± 1.27 | | | |
| **7a** | HEK P | N.A. | N.A. | N.A. | | t-test |
| | ΔGRK | 0.0039 | 37.57 ± 4.96 | | | |
| **7b** | HEK P | N.A | -2.01 ± 3.13 | N.A | -7.95 ± 0.096 | Extra sum-of-squares F test |
| | ΔGRK | <0.0001 | 37.39 ± 3.70 | 0.0055 | -7.68 ± 0.17 | |
| **7c** | HEK P | N.A. | -9.20 ± 18.31 | N.A. | | t-test |
| | ΔGRK | 0.0026 | 119.61 ± 24.80 | | | |
| **7d** | HEK P | N.A | 0.43 ± 3.00 | N.A | -8.67 ± 0.15 | Extra sum-of-squares F test |
| | ΔGRK | <0.0001 | 68.20 ± 4.35 | 0.16 | -8.19 ± 0.65 | |
| **S3** | CCR9 WT | N.A | N.A | N.A. | | t-test |
| | CCR9 ΔCT | 0,029 | 83,39 ± 5,00 | | | |
| **S5b** | HEK P | N.A. | 0.242 ± 0.0214 | N.A. | | t-test |
| | ΔGRK | 0.0007 | 1.22 ± 0.0679 | | | |
| **S8** | CCR9 WT | N.A | N.A | N.A | | Browns-Forsythe & Welch ANOVA |
| | CCR9 C1 | 0.014 | 56.83 ± 8.84 | | | |
| | CCR9 C2 | 0.0006 | 26.83 ± 3.12 | | | |
| | CCR9 C1+C2 | 0.0056 | 24.27 ± 9.84 | | | |
| **S9b** | CCR9 WT | N.A. | 0.46 ± 0.26 | N.A. | | |

| | | | | | |
|---|---|---|---|---|---|
| | CCR9 C1 | 0.010 | 1.54 ± 0.10 | | Browns-Forsythe & Welch ANOVA |
| | CCR9 C2 | 0.75 | 0.38 ± 0.3 | | |
| | CCR9 C1+C2 | 0.0099 | 1.82 ± 0.38 | | |
| **S9c** | CCR9 WT | N.A. | N.A. | N.A. | Browns-Forsythe & Welch ANOVA |
| | CCR9 C1 | 0.27 | 72.04 ± 32.04 | | |
| | CCR9 C2 | 0.0079 | 27.04 ± 11.30 | | |
| | CCR9 C1+C2 | 0.026 | 48.58 ± 14.67 | | |
